# Genomic data support *Betula halophila* and *Betula microphylla* as one species and reveal unidirectional introgression from *Betula pendula* to *Betula microphylla*

**DOI:** 10.1101/2022.10.16.512449

**Authors:** Junyi Ding, Donglai Hua, Linmei Yao, Nian Wang

## Abstract

Conservation of rare species faces challenges arising from uncertainties in species recognition, interspecific gene flow and global climate change. *Betula microphylla* and *Betula halophila* are endangered species in Xinjiang province, Northwest China, where they occur with the abundant *Betula pendula*. The species status of *B. halophila* remains dubious. The extent of gene flow between B. *microphylla* and *B. pendula* remain unexplored. Here, we first resolve the identity of *B. halophila* and then assess the extent of gene flow between *B. microphylla* and *B. pendula* using restriction-site associated DNA sequencing (RADseq). We sequenced 40 *B. pendula* individuals, 40 *B. microphylla* individuals, one *B. halophila* individual and seven *B. tianshanica* individuals. Our molecular analyses show that *B. halophila* and *B. microphylla* refer to the same species. STRUCTURE analyses show unidirectional genetic admixture from *B. pendula* to *B. microphylla*. The ABBA-BABA test indicates that the genetic admixture reflects introgression rather than incomplete lineage sorting. Furthermore, we identified 233 functional genes within the introgressed regions with eight genes related to salt-tolerance, suggesting the possibility of potential adaptive introgression. Our study shows an urgent need to conserve the genetically pure populations of *B. microphylla* and to shift conservation efforts from *B. halophila* to *B. microphylla*. In addition, ex-situ conservation of B. microphylla and conservation strategies to avoid genetic swamping by *B. pendula* and *B. tianshanica* should be implemented.

## Introduction

Interspecific hybridization commonly occurs in nature with ∼25% plant species and 10% animal species estimated to hybridize naturally (Mallet, 2007). It can be facilitated by global climate change via breakdown of geographical isolation or erosion of ecological isolation barriers (Garroway et al., 2010; Vallejo-Marín & Hiscock, 2016). Hybridization is often thought as a mean to increase adaptive potentials via transferring beneficial alleles (Hedrick, 2013; Hamilton & Miller, 2016) (Rieseberg, 1997; Abbott et al., 2013) or generating transgressive traits (Pereira et al. 2013; Seehausen 2013), which may enable species adapt to future climate. However, hybridization may drive rare species to extinction through genetic or demographic swamping (Allendorf et al., 2004; Edmands, 2007; Todesco et al., 2016). Understanding the extent of interspecific hybridization would guide species conservation. Whether to conserve hybridized populations remain debated with traditional views suggesting that hybrids should be eradicated (O’Brien & Mayr, 1991; Jackiw et al., 2015). However, an increasing number of studies advocate to conserve hybridized populations due to the evolutionary significance that hybridization has played (Allendorf et al., 2001; Chan et al., 2019). Second, hybridization has been reported to be associated with environmental disturbances (Crispo et al., 2011; Ortego et al., 2017; Grabenstein & Taylor, 2018). Climate change or human activities may indirectly threaten rare species via facilitating interspecific hybridization.

Additional challenges facing conservation biology arise from uncertainties in species taxonomy (Ennos et al., 2005; Ennos et al., 2012; Garmendia et al., 2022). A rare species may be misidentified and conservation efforts may not be timely allocated (Joppa et al., 2011; Vogel Ely et al., 2017). Alternatively, some populations of a common species may be treated as a rare species, wasting conservation efforts (Milne et al., 2009; Garmendia et al., 2022).

The genus *Betula* (birch) is such a case in point. *Betula* contains approximately 65 trees or shrubs species, which have a wide distribution in the Northern Hemisphere (Ashburner & McAllister, 2016). Some *Betula* species are pioneers and grow in forest gaps (Osumi & Sakurai, 1997; Fischer et al., 2002). *Betula* species has a tough taxonomy with two subgenera (*Aspera and Betula*) established recently based on phylogenomic data (Wang et al., 2021). Introgressive hybridization, polyploidization and considerable morphological variation complicated species delimitation (Nagamitsu et al., 2004; Palme et al., 2004; Thomson et al., 2015; Wang et al., 2016; Hu et al., 2019; Wang et al., 2021). Hybridization and introgression often occur between *Betula* species with different ploidy levels, such as hybridization between *B. nana* and *B. pubescens* with introgression asymmetrically occurred from the former to the latter (Zohren et al., 2016; Tsuda et al., 2017; Maslov et al., 2019; Rousi et al., 2019). Habitat conditions (Eriksson & Jonsson, 1986; Bona et al., 2018), long distance transport of pollen (Rousi et al., 2019) and biogeographical history (Tsuda et al., 2017) can facilitate hybridization between *Betula* species. *Betula* includes several endangered species, which are not only vulnerable to future climate change but also subject to hybridization with other closely-related species.

In the present study, we focus on *B. microphylla, B. halophila* and *B. pendula* in Xinjiang province, Northwest China. *Betula microphylla* (2n = 56) is a tree species thriving in wetland (Ashburner and McAllister, 2016). In past decades, *B. microphylla* has undergone dramatic decrease due to human activities and some wild populations have become extinct (Koropachinskii, 2013). *Betula halophila* is salt-tolerant and has been listed as an endangered species with a second-class level in China (Wang et al., 2003; Guo & Zang, 2013). *Betula halophila* was firstly discovered in the Altay region in 1956 and has gone extinct from its natural habitats. Considerable efforts have been made to conserve *B. halophila*. For example, *B. halophila* seedlings have been successfully planted in Weizishan Nature Reserve in Yantai city and Changdao Nature Reserve in Qingdao city, Shandong province. However, the species status of B. halophila remains dubious and it has been considered as a variety of *B. microphylla* (Li & Skvortsov, 1999). The close relationship between *B. halophila* and *B. microphylla* was supported by ITS-based phylogenetic analysis (Wang et al., 2016). In contrast to the narrowly distributed *B. microphylla, B. pendula* commonly occurs in the northern Xinjiang. Our previous study shows that *B. microphylla* has genetic materials from its allopatric relative *B. tianshanica*, with the southern *B. microphylla* populations having a much higher degree of genetic admixture than the northernmost populations (Ding et al., 2021). Given abundant *B. pendula* individuals in northern Xinjiang, we would expect gene flow between *B. pendula* and *B. microphylla*. Indeed, hybrids between *B. microphylla* and *B. pendula* have been reported based on morphological characters (Koropachinskii, 2013).

In the present study, we first test the hypothesis that *B. halophila* and *B. microphylla* refer to the same species. Then we investigate if gene flow occurs between *B. pendula* and *B. microphylla* using restriction-site associated DNA sequencing (RADseq). To this end, we genotyped 88 samples, representing one *B. halophila*, 40 *B. microphylla*, 40 *B. pendula* and seven *B. tianshanica* using RADseq. Two previous sequenced individuals, representing each of *B. microphylla* and *B. halophila* were also included for population structure and phylogenomic analyses.

## Materials and methods

### DNA extraction and sequencing

Samples used for RADseq were chosen from our collected birches in 2018. Samples include *B. pendula* from eight populations, *B. microphylla* from five populations, *B. halophila* from one population and *B. tianshanica* from one population in NW China. The *B. halophila* population was artificially restored to the site where it was first discovered. Collected samples within each population were separated by 20 m. A total of 88 individuals were sequenced in the present study, including 40 *B. pendula*, 40 *B. microphylla*, seven *B. tianshanica* and one *B. halophila*. Total genomic DNA from cambial tissues was isolated using modified 2x cetyltrimethylammonium bromide (CTAB) method following the protocol of Wang et al. (2013). All individuals were subjected to RAD sequencing using the restriction enzyme PstI and sequenced on an Illumina HiSeq 2500 platform, with paired-end 150 bp sequencing. Construction of RAD libraries, sequencing, adaptors trimming and de-multiplexing were performed at Personalbio company, China.

### Read mapping, variant calling and hard filtering

Bases with a quality of below 20 within the sliding-window of 5 bp were trimmed from the start or the end of reads using Trimmomatic v0.39 (Bolger et al., 2014). After trimming, reads shorter than 40 bp and unpaired reads were discarded. Reads passing quality control were mapped to the *B. pendula* reference genome (Salojärvi et al., 2017) using the BWA-MEM algorithm with default parameters, as implemented in BWA v0.7.17-r1188 (Li & Durbin, 2009). All subsequent analyses, such as marking duplicates, variants calling, GVCF files merging and joint genotyping, were performed in GATK v4.1.8.1 (McKenna et al., 2010; DePristo et al., 2011). Two datasets namely dataset 1 (hereafter D1) and dataset 2 (hereafter D2) were created with D1 treating the tetraploid individuals as diploids to create a pseudo-diploid dataset and with D2 a mixed ploidy dataset as described by Zohren et al. (2016). Both datasets include RAD data of the 40 *B. pendula* individuals, 40 *B. microphylla* individuals, one *B. halophila* and each of the previously sequenced *B. microphylla* and *B. halophila*. To obtain high quality SNPs, several hard filtering steps were employed using GATK and BCFtools v1.10.2 (Li, 2011): (a) we only include biallelic SNPs with MQ greater than 40.0, QUAL greater than 30.0, QD greater than 2.0, SOR below than 3.0, FS below than 60.0, ExcessHet below than 54.69, MQRankSum greater than −12.5, DP between 5 and 200, ReadPosRankSum greater than −8.0 (b) we removed all SNPs with minor allele frequency (MAF) <= 0.01 and r^2^ > 0.5 within a 50 kb window to reduce linkage disequilibrium; (c) no missing genotypes were allowed.

### Population structure analysis

D1 and D2 were analyzed in STRUCTURE v2.3.4 (Pritchard et al., 2000) to infer the optimal number of genetic clusters. To reduce computational costs, we randomly selected 50,000 SNPs sites from each dataset. The optimal K value was set from 1 to 5 with the admixture model and correlated allele frequencies. For each K, five independent runs were performed with a burn-in of 100,000 and MCMC replications of 100,000. The optimal value of clusters was estimated using STRUCTURE HARVESTER v0.6.94 (Earl & vonHoldt, 2012) according to the method described by Evanno et al. (2005). Replicate runs were grouped based on a symmetrical similarity coefficient of >0.9 in CLUMPP v1.1 (Jakobsson & Rosenberg, 2007) and were visualized in DISTRUCT v1.1 (Rosenberg, 2004). In order to compare the admixture proportions yielded from STRUCTURE analyses of two genotyping pipelines, a Mann–Whitney U-test was conducted using “wilcox.test” function in R v4.0.3 (R Core Team, 2020). In addition, to summarize genetic clustering patterns, a principal component analysis (PCA) was performed with the “adegenet” R package v2.1.3 (Jombart, 2008).

### Patterns of introgression

The ABBA-BABA test was used to detect signals of introgression from either *B. pendula* or *B. tianshanica* into *B. microphylla*. A total of 51 individuals were randomly selected for ABBA-BABA test, representing 21 *B. pendula*, 22 *B. microphylla*, seven *B. tianshanica* and an outgroup *B. schimidtii* from Wang et al. (2021). In a phylogeny with topology (((P1,P2))P3)O), an excess of ABBA pattern is expected if gene flow has occurred between P2 and P3, provided that P1 and P3 do not or just limitedly exchange genes. Patterson’s D-statistic was used to detect the asymmetries in the frequency of shared and derived variants (Green et al., 2010; Durand et al., 2011). The D and f4-ratio statistic were calculated using function “Dtrios” in Dsuite 0.4-r38 (Malinsky et al., 2021). A block jack-knifing was performed to obtain an associated p value for the test of whether D statistic deviates significantly from zero. Z score was calculated for the following topologies: (*B. schimidtii*, (*B. pendula*, (*B. microphylla, B. tianshanica*))), (*B. schimidtii*, (*B. pendula*, (*B. microphylla, B. microphylla*))) and (*B. schimidtii*, (*B. tianshanica*, (*B. microphylla, B. microphylla*))). The median Z score larger than 3.10 (equal to a p value < 0.001 of D statistic) was regarded as significant signals of introgression.

### Identification of introgressed regions

Putative introgressed regions were identified using f_DM_ statistics, a powerful method to detect introgressed signal in short genomic regions (Malinsky et al., 2015). The “Dinvestigate” function was used to calculate f_DM_ statistics on a sliding non-overlapping window size of 50 SNPs incremented by 25 SNPs across the genome. A candidate window with the highest 1 % of f_DM_ values was thought to harbor introgressed loci (Etherington et al., 2022). The putative introgressed regions were defined as 10,000 bp flanking each side of SNPs within candidate windows and overlapping regions were merged. In order to understand the biological functions of the introgressed regions, Gene Ontology (GO) analysis was carried out for the putative introgressed regions that overlapped with the annotated B. pendula genes. Enrichment analysis was performed using agriGO v2.0 (Tian et al., 2017) based on Fisher’s exact test and GO terms with a FDR cutoff of⍰0.001 were treated as significant in this study.

### Inference of ploidy level

We inferred the ploidy of *B. halophila* and *B. microphylla* in this study following the method in Zohren et al. (2016).

### Phylogenetic analyses

To determine the phylogenetic position of *B. halophila*, we created a dataset of 30 samples, including 23 *Betula* taxa from a previous study (Wang et al., 2021), three *B. microphylla*, three *B. tianshanica* and one *B. halophila* generated in this study. SNPs with a missing rate greater than 50% were removed. A phylogenetic tree of the concatenated SNPs was constructed using RAxML v8.2.12 (Stamatakis, 2006) with the GTR+GAMMA model. Alnus inokumae was selected as the outgroup. Support values were calculated based on 100 bootstraps.

## RESULTS

### RAD-seq and SNP calling

An average of 16,151,364 reads was yielded per individual with 15,550,747 retained for each individual after quality filtering. Between 10,590,234 and 26,541,500 reads were mapped to the reference (Table S1). After filtering, 91,902 and 75,639 SNPs were retained in D1 in D2, respectively. Raw restriction-site associated DNA sequencing data generated for this study have been deposited in the NCBI Sequence Read Archive (SRA) under accession number PRJNA883437.

### Population genetic structure

The STRUCTURE results based on D1 and D2 are similar (p = 0.742, Mann-Whitney U test) and only the result of D1 is shown in the main text (Figure 1; Figures S1-2). The optimal number of genetic clusters is two, corresponding to *B. pendula* and *B. microphylla*/*B. halophila* (Figure S1). When K =2, *B. microphylla* from population JMN revealed obvious admixture from *B. pendula* (13.74% ± 0.03) (Figure 1A). The two *B. halophila* individuals have similar genetic composition with *B. microphylla* and showed admixture from *B. pendula* (Figure 1AC). Higher K values (3 and 4) did not subdivide *B. microphylla* populations (Figure S2).

**Figure 1.**
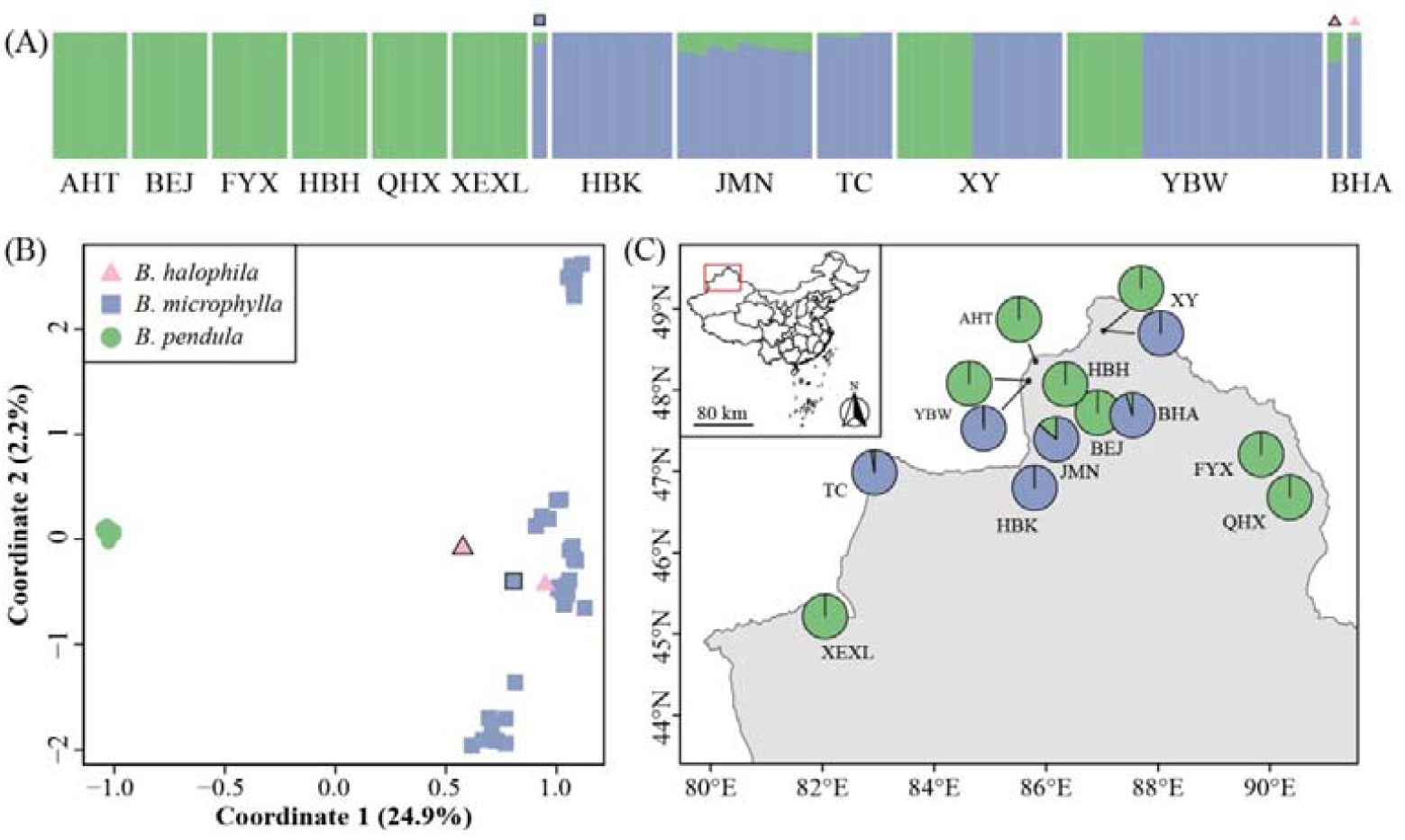
(A) STRUCTURE analysis of *B. microphylla, B. pendula* and *B. halophila* at K=2 based on the 50,000 pseudo-diploid SNPs. *Betula microphylla* and *B. halophila* samples from Wang et al. (2021) are shown by blue square and pink triangle with solid outer lines, respectively. (B) Principal component analysis (PCA) of *B. microphylla, B. pendula* and *B. halophila* based on the 91,902 pseudo-diploid SNPs. (C) A map showing the genetic composition of *B. microphylla* and *B. pendula* populations. The sampling locality is represented by the centre of pie charts or dots which were connected to pie charts by a straight line.

Consistent with STRUCTURE analyses, the PCA result based on D1 showed that *B. pendula* was clearly separated from *B. microphylla* whereas *B. halophila* and *B. microphylla* were grouped together on PC1, which explained 24.9% of variation. PC2 revealed sub-clusters in *B. microphylla*, but only explained 2.2% of variation (Figure 1B). The PCA results based on D1 and D2 are similar (Figure S3).

### Introgression

The ABBA-BABA test suggests a significant signal of introgression from *B. pendula* to *B. microphylla* (median D-statistic = 0.0976, median Z-score = 5.20) and from *B. tianshanica* to *B. microphylla* (median D-statistic = 0.0424, median Z-score = 4.72) (Table 1, Figure 2A). f4-ratio statistic shows that 14.7% of *B. microphylla* genome originates from *B. pendula* (Figure 2B). We observed the highest mean Z-scores in JMN population (Table 1) among the trio of *B. microphylla* and *B. pendula*, where the STRUCTURE plots suggested some allele sharing.

**Table 1.** The results of ABBA-BABA test.

| Topology<sup>1</sup> | Population | Mean<br>Z score | Mean D<br>score | Mean<br>BBAA site | Mean<br>ABBA site | Mean BABA<br>site | Number<br>of trios |
| --- | --- | --- | --- | --- | --- | --- | --- |
| ((M, M), P) |  |  |  |  |  |  |  |
|  | HBK | 0.3 | 0.0025 | 11762 | 6355 | 6323 | 12 |
|  | JMN | 9.5 | 0.18 | 10080 | 8426 | 5834 | 2016 |
|  | TC | 2.8 | 0.052 | 11363 | 6955 | 6260 | 715 |
|  | XY | 1.5 | 0.02 | 11745 | 6427 | 6173 | 428 |
|  | YBW | 3.1 | 0.043 | 11726 | 6698 | 6147 | 840 |
| ((M, M), T) |  |  |  |  |  |  |  |
|  | HBK | 4.8 | 0.039 | 9060 | 8711 | 8060 | 372 |
|  | JMN | - | - | - | - | - | 0 |
|  | TC | 3.2 | 0.03 | 9184 | 8641 | 8144 | 214 |
|  | XY | 4 | 0.04 | 9290 | 8514 | 7854 | 228 |
|  | YBW | 4.7 | 0.048 | 9312 | 8666 | 7866 | 195 |

<sup>1</sup>Letters T, M and P represent *B. tianshanica*, *B. microphylla* and *B. pendula*, respectively.

**Figure 2.**
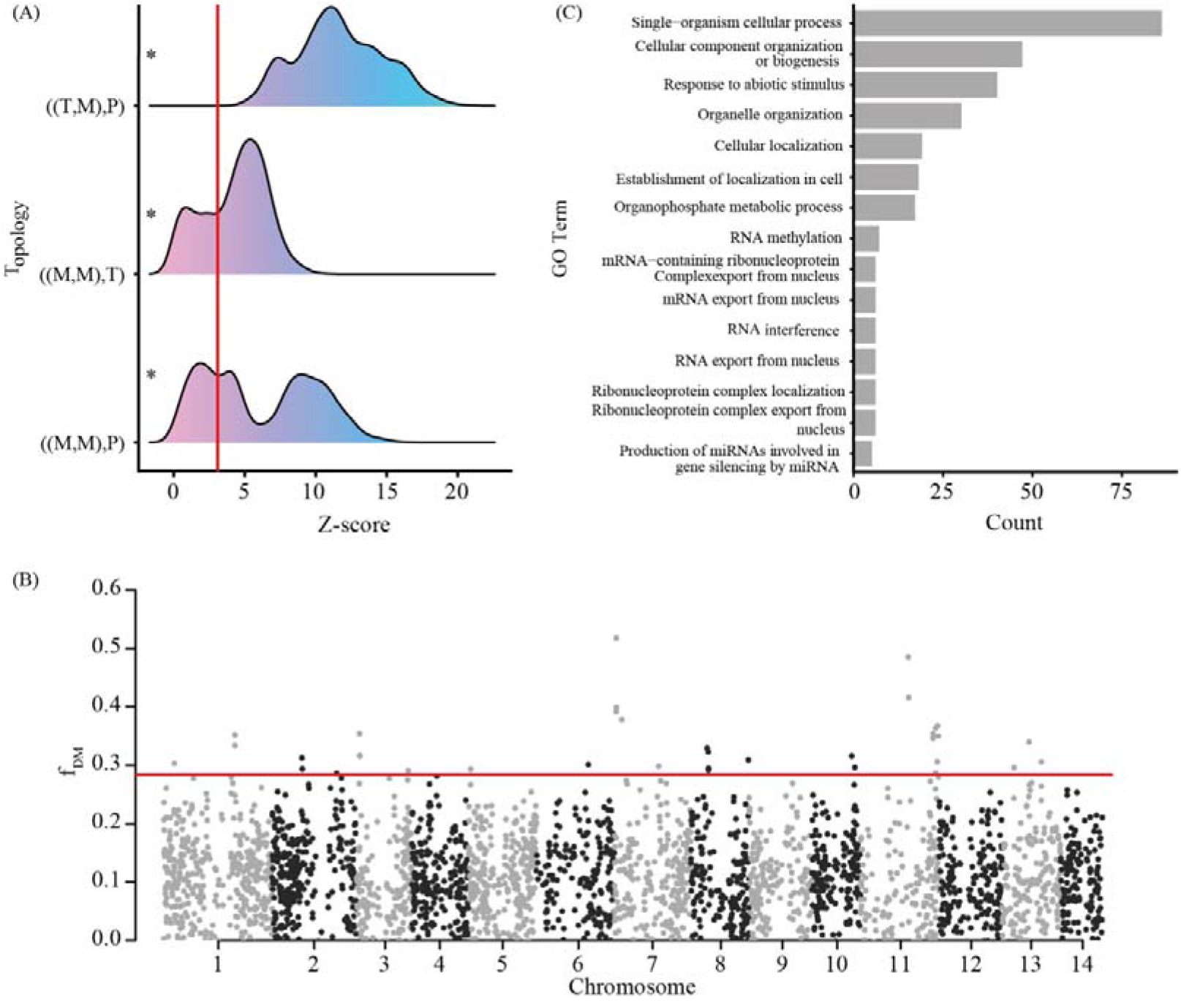
Pattern of introgression between Betula species. (A) Z scores for “ABBA-BABA” introgression tests. Distributions are over combinations of individuals corresponding to each reasonable topology. Topologies are listed as ((P1, P2), P3), with abbreviations T = *B. tianshanica*, M = *B. microphylla*, P = *B. pendula*. Asterisks were used to denote significant introgression topologies, which are indicated by a median Z score > 3.10. (B) Manhattan plot showing the putative introgressed regions between *B. pendula* and *B. microphylla*. (C) GO enrichment analysis results (FDR < 0.001) of the 233 genes spanned by the 81 highly introgressed regions identified with fDM statistic.

After merged overlapping windows, we found 81 highly introgressed regions (f_DM_ 0.29 or above) in *B. microphylla* genome, which originated from *B. pendula*. Within these regions, there are a total of 234 genes, with an average of 2.9 genes per introgressed region. One un-annotatable gene based on *B. pendula* gene models was excluded for enrichment analysis, resulting in 233 genes. Single-organism cellular process (count: 86), cellular component organization or biogenesis (count: 47), and response to abiotic stimulus (count: 40) are the top three GO terms for the enriched genes (Figure 2C). Among these, eight genes were responsive to salt stress (Table S2).

### Ploidy level inference

Allele ratios at heterozygous sites of *B. halophila, B. microphylla* (including the introgressed individuals), *B. tianshanica* showed peaks close to 0.25, 0.50 and 0.75, indicating that they are allotetraploid (Figure S4).

### Phylogenetic position of B. halophila

The phylogenetic tree based on 1,384,918 SNPs showed that *B. halophila, B. microphylla, B. tianshanica* and *B. humilis* formed a monophyletic clade with 100% support (Figure S5).

## Discussion

### Betula halophila and B. microphylla refer to the same species

*Betula halophila* was first discovered in 1956 and was listed as a critically endangered species (Wang et al., 2003; Guo & Zang, 2013). However, *B. halophila* was dubiously regarded as *B. microphylla* (Ashburner & McAllister, 2016). Here, our genetic analyses support *B. halophila* and *B. microphylla* as one species. We found *B. halophila* to be *B. microphylla* genetically with genetic admixture from *B. pendula*. We showed that *B. halophila* is a tetraploid, consistent with the ploidy of *B. microphylla* (Ashburner & McAllister, 2016; Wang et al., 2016). Geographically, *B. halophila* is within the range of *B. microphylla*. Taken these together, we suggest that *B. halophila* should be treated as *B. microphylla*. We found *B. halophila, B. microphylla* and *B. tianshanica* formed a monophyletic clade with *B. humilis*, supporting *B. humilis* as their shared diploid progenitor (Wang et al., 2021).

### Introgression from *B. pendula* to *B. microphylla*

We observed a high level of genetic admixture from *B. pendula* to *B. microphylla* and ascribed the genetic admixture to introgressive hybridization rather than incomplete lineage sorting (ILS). First, ABBA-BABA test shows significant signals of introgression, which would not be expected under the scenario of ILS. Second, the introgressed regions are localized in several chromosomes rather than randomly distributed across the genome, indicating that ILS is not a main cause to the observed patterns of admixture (Taylor et al., 2020). The occurrence of introgression between *B. pendula* and *B. microphylla* suggests that a difference in ploidy cannot prevent gene flow as demonstrated in several other plant species (Sutherland & Galloway, 2017; Wang et al., 2020). As expected, the direction of introgression is from diploid to tetraploid fitting with the hypothesis proposed by Stebbins (1971) and is consistent with previous studies showing unidirectional introgression from diploid to tetraploid species, such as from *Betula* nana to *B. pubescens* (Zohren et al., 2016) and from *Miscanthus sinensis* to *M. sacchariflorus* (Clark et al., 2018). Our study reveals the absence of intorgression in sympatric populations of *B. microphylla* and *B. pendula* (YBW and XY), suggesting species integrity was maintained there (Figure 2A). The absence of introgression between sympatric species has been indicated by a previous studies investigating spatial patterns of hybridization (Hasselman et al., 2014). In contrast, our study reveals that population JMN was a hybrid swarm and all individuals there were deeply introgressed, reflecting several generations of backcrossing. This suggests that the postzygotic reproductive isolation may have broken down and there may not be selection against hybrids in population JMN. We are not clear why isolation barriers between *B. microphylla* and *B. platyphylla* varied spatially. Anthropogenic disturbances may play a role in explaining such a difference, as demonstrated across many other taxa (Hubbs, 1955; Guo, 2014; Beddows & Rose, 2018). We found that the introgressed regions from *B. pendula* to *B. microphylla* harbor functional genes involved in cellular process, cellular component organization and biogenesis and response to abiotic stimuli (Figure 2C). This suggests that introgression from *B. pendula* to *B. microphylla* may be adaptive. Interestingly, we noted that *B. halophila* was also introgressed from *B. pendula*. The fact that *B. halophila* was salt-tolerant suggests that introgression from *B. pendula* may be adaptive (Zhang et al., 2009; Shao et al., 2018). However, we did not note any studies comparing salt-tolerance between introgressed *B. microphylla* from *B. pendula* and purebred *B. microphylla*. Further research is needed to test this hypothesis.

### Conservation practices

Ennos et al. (2012) has developed a process-based approach to protect taxonomically complex groups by conserving evolutionary processes, such as hybridization and introgression. We support to implement the process-based approach to conserve *B. microphylla* as suggested previously (Ding et al., 2021). We unexpectedly found that *B. microphylla* from population JMN was deeply introgressed and the introgressed regions contain genes involving in response to abiotic stimuli. Therefore, we advocate conserving population JMN as *B. microphylla* there may adapt to future climate change. We found that populations XY and YBW represent genetically purebred *B. microphylla*. Conservation efforts should be allocated to conserve the two populations to maintain the genetic purity of *B. microphylla*. Meanwhile, we suggest that efforts to conserve *B. halophila* should be shifted toward conserving *B. microphylla* as our results clearly indicate that *B. halophila* is introgressed *B. microphylla* from *B. pendula*. Furthermore, we put an emphasis on protecting habitats of *B. microphylla*. On one hand, anthropogenic disturbance, such as crop plantation, has directly driven some *B. microphylla* populations extinct by habitat alteration. On another hand, anthropogenic disturbance has been reported to be associated with hybridization (Anderson, 1948; Ortego et al., 2017; Grabenstein & Taylor, 2018). Although we did not detect introgression from *B. pendula* to *B. microphylla* in populations XY and YBW, gene flow from *B. pendula* remains a potential threat. Given the extremely abundant *B. pendula* individuals in populations XY and YBW, genetic swamping of B. microphylla may be possible in the future once the current reproductive barriers were broken down. Lastly, we advocate conserving *B. microphylla* ex situ as this species has been shown to thrive in costal areas in Shandong province (Huang et al., 2020).

## Acknowledgements

This work was funded by the National Natural Science Foundation of China (31770230), the Program for Introduction and Cultivation of Young Scholars in Universities in Shandong Province and the Outstanding Young Scholars funded by the State Forestry and Grassland Administration, China.

