## Supplemental tables and figures for "Genomic data support *Betula halophila* and *Betula microphylla* as one species and reveal unidirectional introgression from *Betula pendula* to *Betula microphylla*"

**Table S1** Detailed information on samples used for RADseq and RADseq results.

**Table S2** Detailed information on introgressed genes responded to salt stress.

**Figure S1** The best number of clusters inferred using the “Evanno test” method based on (a) pseudo-diploid dataset and (b) mixed ploidy dataset.

**Figure S2** STRUCTURE results at K values from 2 to 4 based on (a) pseudo-diploid dataset and (b) mixed ploidy dataset at randomly selected 50,000 SNPs sites. Square and triangle with solid line represent previously sequenced samples in Wang et al. (2021).

**Figure S3** The result of principal component analysis (PCoA) based on mixed ploidy dataset at 75,639 SNPs. Square and triangle with solid line represent samples Wang et al. (2021).

**Figure S4** Plots of read count ratios for heterozygous sites covered by at least 30 reads for *B. microphylla*, *B. halophila* and *B. tianshanica*.

**Figure S5** Phylogenetic tree from the maximum-likelihood analysis of *B. halophila*, *B. microphylla* and *B. tianshanica* using the supermatrix approach based on data from 1,384,918 SNPs. The scale bar below indicates the mean number of nucleotide substitutions per site. Bootstrap percentages of 100 % are not shown. The colored indicates the sequenced samples in the present study.

**Table S1** Detailed information on samples used for RADseq and RADseq results.

| Sample ID | Species | Latitude (°N) | Longitude (°E) | Clean reads | Mapped reads | Percentage (%) | Variants |
| --- | --- | --- | --- | --- | --- | --- | --- |
| AHT003 | *B. pendula* | 48.38 | 85.75 | 16,013,526 | 15,049,864 | 93.98 | 495,726 |
| AHT011 | *B. pendula* | 48.38 | 85.75 | 16,144,296 | 14,959,212 | 92.66 | 468,940 |
| AHT012 | *B. pendula* | 48.38 | 85.75 | 13,285,444 | 11,912,530 | 89.67 | 435,639 |
| AHT018 | *B. pendula* | 48.38 | 85.75 | 17,685,410 | 16,331,575 | 92.34 | 476,777 |
| AHT025 | *B. pendula* | 48.38 | 85.75 | 18,738,544 | 17,054,433 | 91.01 | 509,002 |
| BEJ002 | *B. pendula* | 47.73 | 86.92 | 17,048,850 | 16,141,809 | 94.68 | 497,485 |
| BEJ008 | *B. pendula* | 47.73 | 86.92 | 28,038,358 | 26,541,500 | 94.66 | 614,601 |
| BEJ015 | *B. pendula* | 47.73 | 86.92 | 14,000,472 | 13,261,805 | 94.72 | 458,004 |
| BEJ017 | *B. pendula* | 47.73 | 86.92 | 14,534,270 | 13,730,942 | 94.47 | 487,612 |
| BEJ021 | *B. pendula* | 47.73 | 86.92 | 15,632,866 | 14,771,978 | 94.49 | 489,129 |
| FYX002 | *B. pendula* | 47.21 | 89.84 | 12,147,610 | 11,459,144 | 94.33 | 495,455 |
| FYX006 | *B. pendula* | 47.21 | 89.84 | 12,706,606 | 12,083,590 | 95.10 | 485,112 |
| FYX010 | *B. pendula* | 47.21 | 89.84 | 14,667,286 | 13,875,776 | 94.60 | 473,401 |
| FYX012 | *B. pendula* | 47.21 | 89.84 | 13,348,854 | 12,629,437 | 94.61 | 475,455 |
| FYX017 | *B. pendula* | 47.21 | 89.84 | 12,531,022 | 11,934,389 | 95.24 | 432,152 |
| HBH001 | *B. pendula* | 48.07 | 86.34 | 13,604,142 | 11,350,256 | 83.43 | 481,671 |
| HBH005 | *B. pendula* | 48.07 | 86.34 | 12,023,402 | 11,294,854 | 93.94 | 454,361 |
| HBH008 | *B. pendula* | 48.07 | 86.34 | 13,159,156 | 12,327,735 | 93.68 | 456,802 |
| HBH011 | *B. pendula* | 48.07 | 86.34 | 15,273,324 | 14,290,701 | 93.57 | 530,968 |
| HBH020 | *B. pendula* | 48.07 | 86.34 | 18,652,508 | 17,462,461 | 93.62 | 494,034 |
| XY-p001 | *B. pendula* | 48.68 | 87.01 | 12,955,764 | 10,590,234 | 81.74 | 1,201,944 |
| XY-p004 | *B. pendula* | 48.68 | 87.01 | 12,308,666 | 11,602,717 | 94.26 | 456,220 |
| XY-p008 | *B. pendula* | 48.68 | 87.01 | 13,348,996 | 12,627,534 | 94.60 | 452,428 |
| XY-p011 | *B. pendula* | 48.68 | 87.01 | 12,575,892 | 11,873,561 | 94.42 | 448,125 |
| XT-p014 | *B. pendula* | 48.68 | 87.01 | 13,321,206 | 12,589,620 | 94.51 | 459,812 |
| QHX001 | *B. pendula* | 46.68 | 90.35 | 15,359,880 | 14,525,702 | 94.57 | 484,891 |
| QHX003 | *B. pendula* | 46.68 | 90.35 | 14,148,734 | 13,408,111 | 94.77 | 518,564 |
| QHX005 | *B. pendula* | 46.68 | 90.35 | 14,548,214 | 13,764,611 | 94.61 | 461,704 |
| QHX008 | *B. pendula* | 46.68 | 90.35 | 15,318,210 | 14,499,449 | 94.66 | 529,617 |
| QHX014 | *B. pendula* | 46.68 | 90.35 | 13,573,670 | 12,854,869 | 94.70 | 491,464 |
| XEXL003 | *B. pendula* | 45.22 | 82.05 | 18,324,306 | 17,365,987 | 94.77 | 453,760 |
| XEXL009 | *B. pendula* | 45.22 | 82.05 | 15,651,688 | 14,888,663 | 95.13 | 475,472 |
| XEXL011 | *B. pendula* | 45.22 | 82.05 | 15,827,566 | 14,616,262 | 92.35 | 495,788 |
| XEXL018 | *B. pendula* | 45.22 | 82.05 | 13,549,158 | 12,915,107 | 95.32 | 448,514 |
| XEXL022 | *B. pendula* | 45.22 | 82.05 | 17,468,314 | 16,644,172 | 95.28 | 452,400 |
| YBW001 | *B. pendula* | 48.12 | 85.69 | 14,102,734 | 13,283,395 | 94.19 | 477,840 |
| YBW005 | *B. pendula* | 48.12 | 85.69 | 13,345,008 | 12,619,865 | 94.57 | 444,192 |
| YBW014 | *B. pendula* | 48.12 | 85.69 | 13,094,166 | 12,388,013 | 94.61 | 463,026 |
| YBW019 | *B. pendula* | 48.12 | 85.69 | 16,511,342 | 15,605,277 | 94.51 | 490,881 |
| YBW025 | *B. pendula* | 48.12 | 85.69 | 15,278,316 | 14,456,712 | 94.62 | 482,053 |
| HBK005 | *B. microphylla* | 46.78 | 85.80 | 17,163,228 | 16,192,195 | 94.34 | 1,081,934 |
| HBK012 | *B. microphylla* | 46.78 | 85.80 | 15,844,014 | 14,984,064 | 94.57 | 1,071,831 |
| HBK014 | *B. microphylla* | 46.78 | 85.80 | 12,349,336 | 11,666,338 | 94.47 | 1,034,049 |
| HBK019 | *B. microphylla* | 46.78 | 85.80 | 13,965,444 | 13,178,712 | 94.37 | 1,095,601 |
| HBK022 | *B. microphylla* | 46.78 | 85.80 | 15,925,668 | 15,034,717 | 94.41 | 1,081,576 |
| HBK024 | *B. microphylla* | 46.78 | 85.80 | 19,084,840 | 17,985,435 | 94.24 | 1,164,039 |
| HBK030 | *B. microphylla* | 46.78 | 85.80 | 14,057,096 | 13,227,761 | 94.10 | 1,127,766 |
| HBK032 | *B. microphylla* | 46.78 | 85.80 | 25,479,592 | 24,029,877 | 94.31 | 1,369,384 |
| JMN007 | *B. microphylla* | 47.38 | 86.18 | 15,352,808 | 14,528,636 | 94.63 | 1,043,264 |
| JMN013 | *B. microphylla* | 47.38 | 86.18 | 15,288,020 | 14,457,050 | 94.56 | 1,023,634 |
| JMN015 | *B. microphylla* | 47.38 | 86.18 | 15,082,746 | 14,233,148 | 94.37 | 1,029,970 |
| JMN018 | *B. microphylla* | 47.38 | 86.18 | 16,736,904 | 15,846,309 | 94.68 | 1,066,197 |
| JMN021 | *B. microphylla* | 47.38 | 86.18 | 12,348,162 | 11,682,916 | 94.61 | 1,026,496 |
| JMN023 | *B. microphylla* | 47.38 | 86.18 | 14,264,494 | 13,482,537 | 94.52 | 1,008,313 |
| JMN026 | *B. microphylla* | 47.38 | 86.18 | 17,041,062 | 16,102,259 | 94.49 | 1,102,163 |
| JMN028 | *B. microphylla* | 47.38 | 86.18 | 17,090,170 | 16,087,437 | 94.13 | 1,341,964 |
| JMN030 | *B. microphylla* | 47.38 | 86.18 | 17,030,392 | 16,029,172 | 94.12 | 1,106,328 |
| TC002 | *B. microphylla* | 46.97 | 82.93 | 15,153,730 | 14,270,193 | 94.17 | 1,069,552 |
| TC004 | *B. microphylla* | 46.97 | 82.93 | 15,611,514 | 14,653,218 | 93.86 | 1,068,453 |
| TC006 | *B. microphylla* | 46.97 | 82.93 | 17,148,866 | 15,944,341 | 92.98 | 1,066,439 |
| TC008 | *B. microphylla* | 46.97 | 82.93 | 15,334,452 | 14,408,251 | 93.96 | 1,076,446 |
| TC009 | *B. microphylla* | 46.97 | 82.93 | 15,395,932 | 14,119,716 | 91.71 | 1,052,346 |
| XY-m004 | *B. microphylla* | 48.72 | 87.03 | 15,526,082 | 14,660,026 | 94.42 | 1,051,707 |
| XY-m006 | *B. microphylla* | 48.72 | 87.03 | 12,880,804 | 12,073,574 | 93.73 | 1,071,871 |
| XY-m008 | *B. microphylla* | 48.72 | 87.03 | 16,166,158 | 15,254,091 | 94.36 | 1,184,513 |
| XY-m011 | *B. microphylla* | 48.72 | 87.03 | 15,061,838 | 14,159,783 | 94.01 | 1,093,304 |
| XY-m016 | *B. microphylla* | 48.72 | 87.03 | 15,620,550 | 14,761,700 | 94.50 | 1,087,290 |
| XY-m021 | *B. microphylla* | 48.72 | 87.03 | 15,847,798 | 14,978,991 | 94.52 | 1,076,907 |
| YBW012 | *B. microphylla* | 48.12 | 85.69 | 17,214,374 | 16,317,755 | 94.79 | 1,139,348 |
| YBW017 | *B. microphylla* | 48.12 | 85.69 | 17,487,258 | 16,536,557 | 94.56 | 1,119,761 |
| YBW021 | *B. microphylla* | 48.12 | 85.69 | 19,495,754 | 17,042,019 | 87.41 | 1,123,643 |
| YBW027 | *B. microphylla* | 48.12 | 85.69 | 15,914,790 | 15,052,036 | 94.58 | 1,264,416 |
| YBW029 | *B. microphylla* | 48.12 | 85.69 | 17,767,526 | 16,785,736 | 94.47 | 1,193,838 |
| YBW033 | *B. microphylla* | 48.12 | 85.69 | 20,287,158 | 19,163,122 | 94.46 | 1,127,342 |
| YBW040 | *B. microphylla* | 48.12 | 85.69 | 15,950,270 | 15,076,133 | 94.52 | 1,085,272 |
| YBW042 | *B. microphylla* | 48.12 | 85.69 | 15,639,092 | 14,772,666 | 94.46 | 1,115,472 |
| YBW044 | *B. microphylla* | 48.12 | 85.69 | 16,062,324 | 15,196,656 | 94.61 | 1,099,685 |
| YBW047 | *B. microphylla* | 48.12 | 85.69 | 14,064,438 | 13,271,930 | 94.37 | 1,081,344 |
| YBW050 | *B. microphylla* | 48.12 | 85.69 | 17,411,956 | 16,312,555 | 93.69 | 1,167,898 |
| YBW053 | *B. microphylla* | 48.12 | 85.69 | 14,908,904 | 14,096,185 | 94.55 | 1,090,360 |
| SLM001 | *B. tianshanica* | 44.43 | 81.06 | 17,172,254 | 16,203,813 | 94.36 | 1,147,885 |
| SLM008 | *B. tianshanica* | 44.43 | 81.06 | 17,293,884 | 16,322,944 | 94.39 | 1,116,345 |
| SLM010 | *B. tianshanica* | 44.43 | 81.06 | 17,092,610 | 16,146,897 | 94.47 | 1,100,653 |
| SLM013 | *B. tianshanica* | 44.43 | 81.06 | 13,072,868 | 12,360,387 | 94.55 | 1,074,789 |
| SLM015 | *B. tianshanica* | 44.43 | 81.06 | 12,660,466 | 11,929,290 | 94.22 | 1,052,192 |
| SLM018 | *B. tianshanica* | 44.43 | 81.06 | 14,158,712 | 13,393,122 | 94.59 | 1,130,749 |
| SLM022 | *B. tianshanica* | 44.43 | 81.06 | 16,905,552 | 16,007,768 | 94.69 | 1,131,673 |
| BHA001 | *B. halophila* | 47.69 | 87.54 | 14,206,066 | 13,422,902 | 94.49 | 1,073,198 |

**Table S2** Detailed information on introgressed genes responded to salt stress.

| Birch Gene ID | Arabidopsis gene with highest sequence similarity | Annotation of Arabidopsis gene |
| --- | --- | --- |
| Bpev01.c2707.g0004 | AT1G65930 | Isocitrate dehydrogenase |
| Bpev01.c0616.g0013 | AT2G36530 | Enolase LENGTH 444 |
| Bpev01.c0356.g0015 | AT4G19810 | Chitinase-3-like protein 2 |
| Bpev01.c0356.g0016 | AT4G19810 | Chitinase-3-like protein 1 |
| Bpev01.c1130.g0010 | AT4G39090 | Cathepsin B-like cysteine proteinase |
| Bpev01.c1394.g0002 | AT5G12380 | annexin 8 LENGTH 316 |
| Bpev01.c0268.g0017 | AT5G17920 | 5-methyltetrahydropteroyltriglutamate--homocysteine methyltransferase |
| Bpev01.c0429.g0008 | AT5G17920 | 5-methyltetrahydropteroyltriglutamate--homocysteine methyltransferase |

**Figure S1** The best number of clusters inferred using the “Evanno test” method based on (a) pseudo-diploid dataset and (b) mixed ploidy dataset.

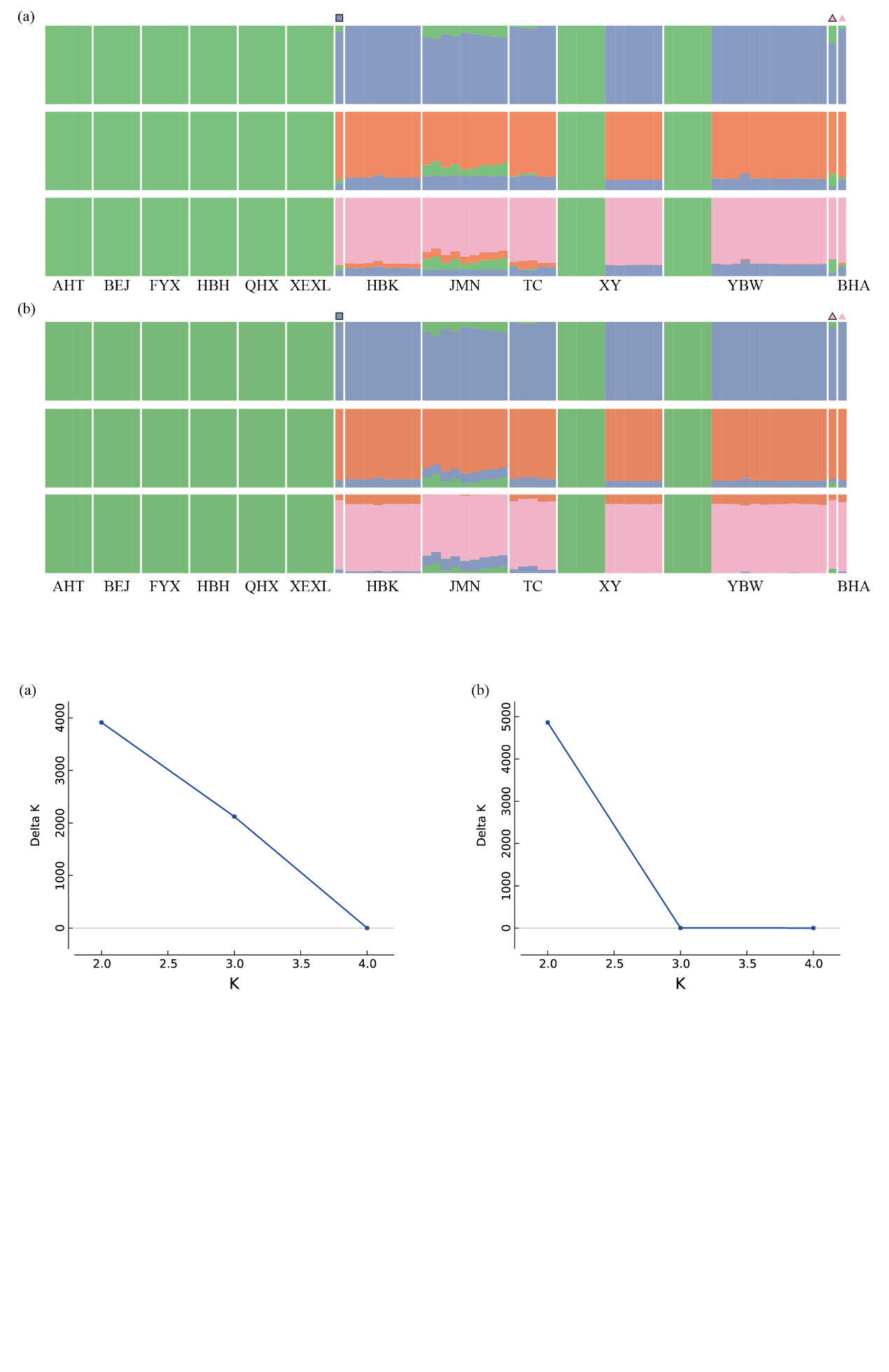

**Figure S2** STRUCTURE results at K values from 2 to 4 based on (a) pseudo-diploid dataset and (b) mixed ploidy dataset at randomly selected 50,000 SNPs sites. Square and triangle with solid line represent previously sequenced samples in Wang et al. (2021).

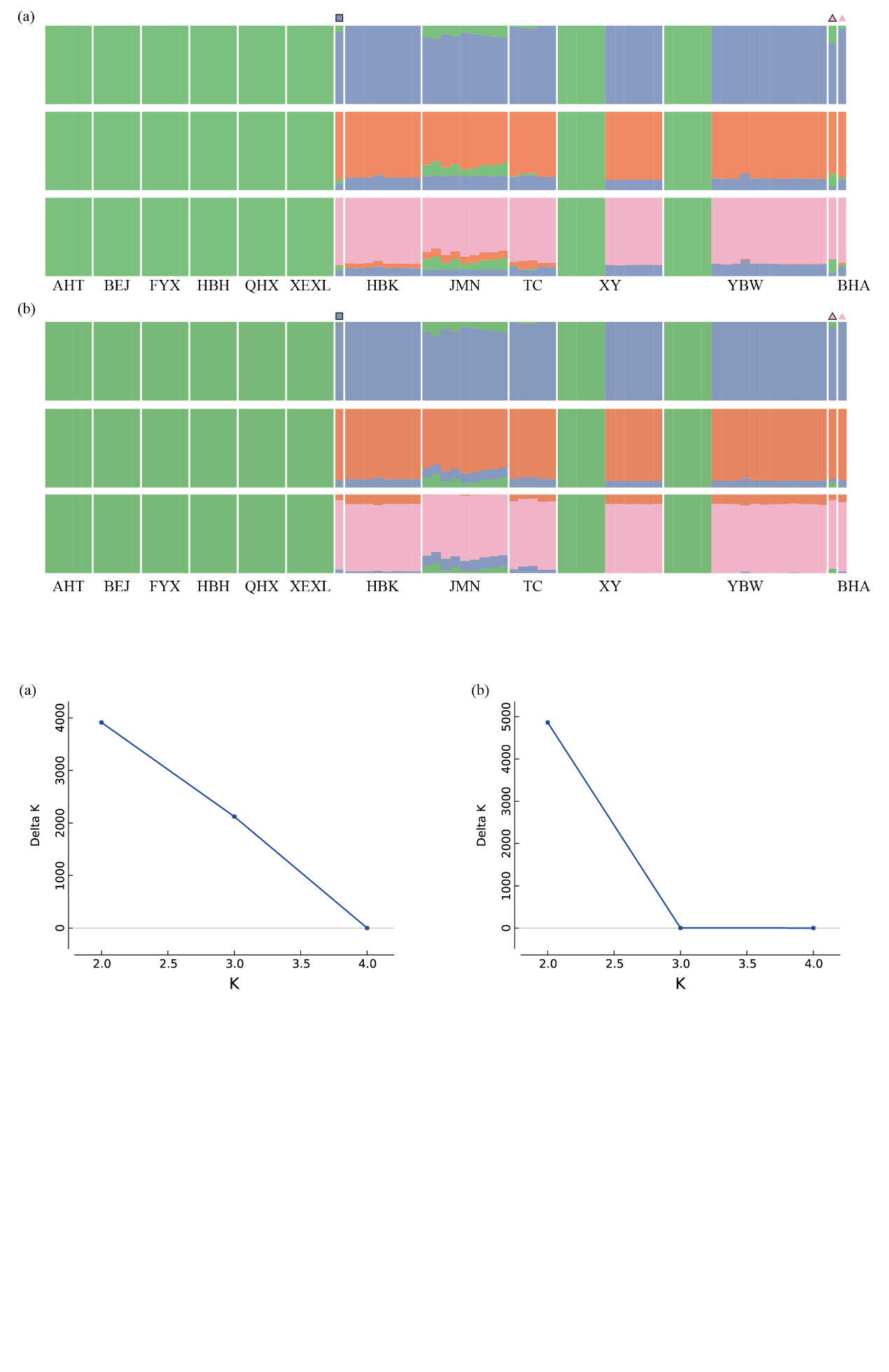

**Figure S3** The result of principal component analysis (PCoA) based on mixed ploidy dataset at 75,639 SNPs. Square and triangle with solid line represent samples Wang et al. (2021).

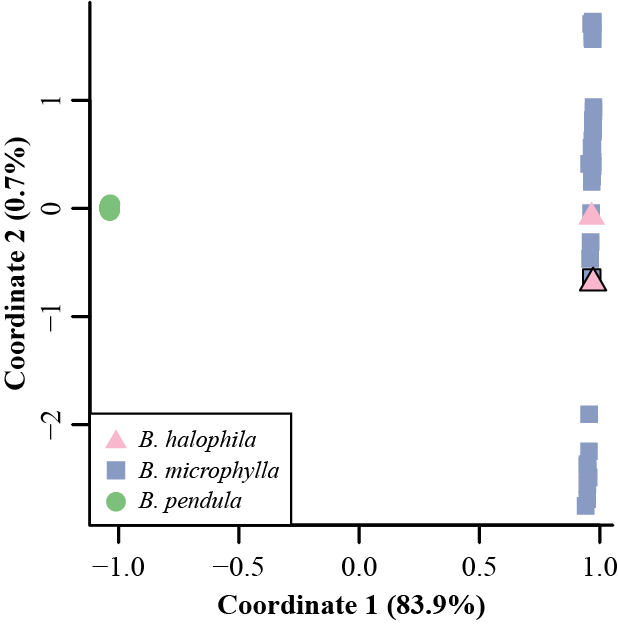

**Figure S4** Plots of read count ratios for heterozygous sites covered by at least 30 reads for *B. microphylla*, *B. halophila* and *B. tianshanica*.

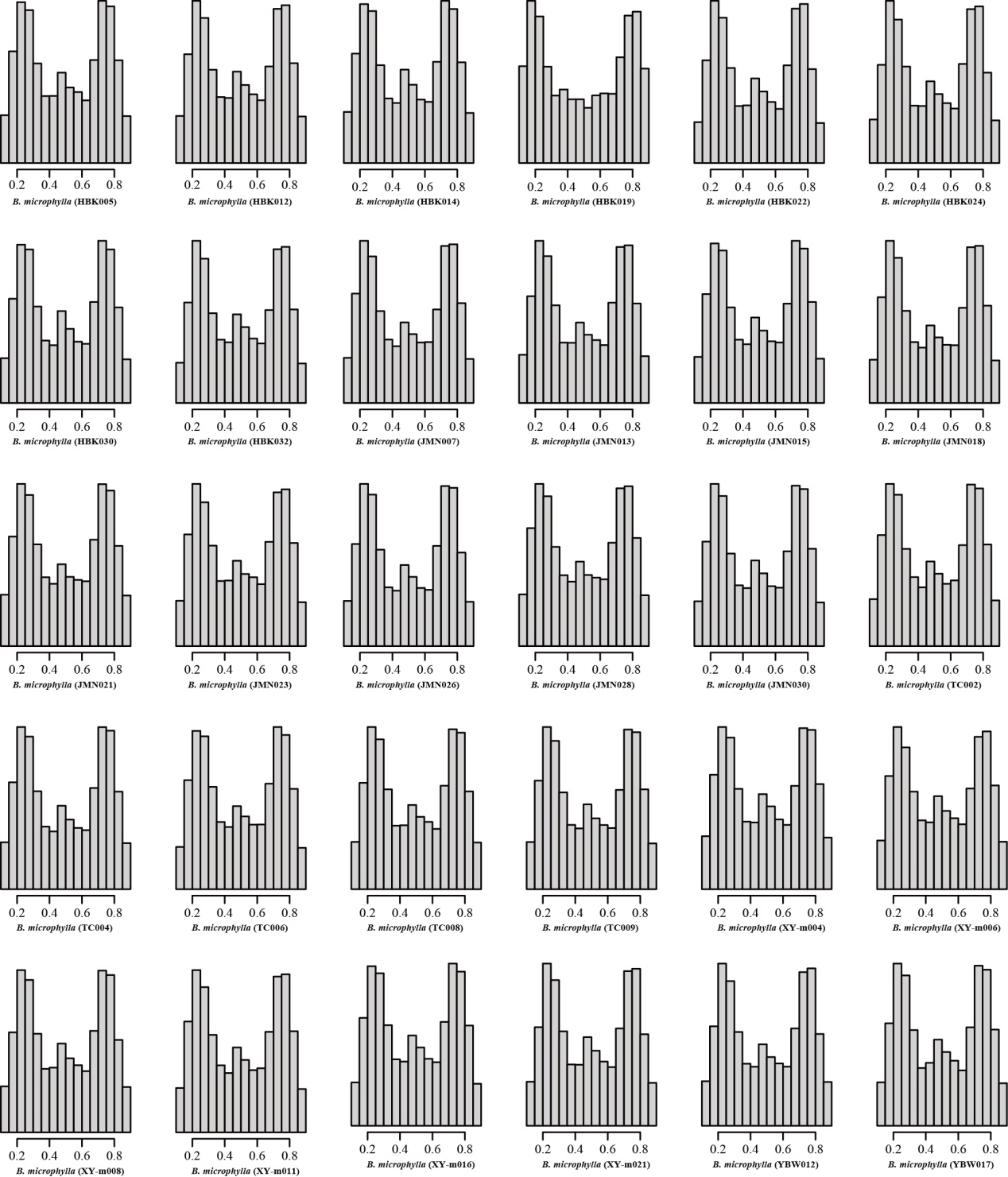

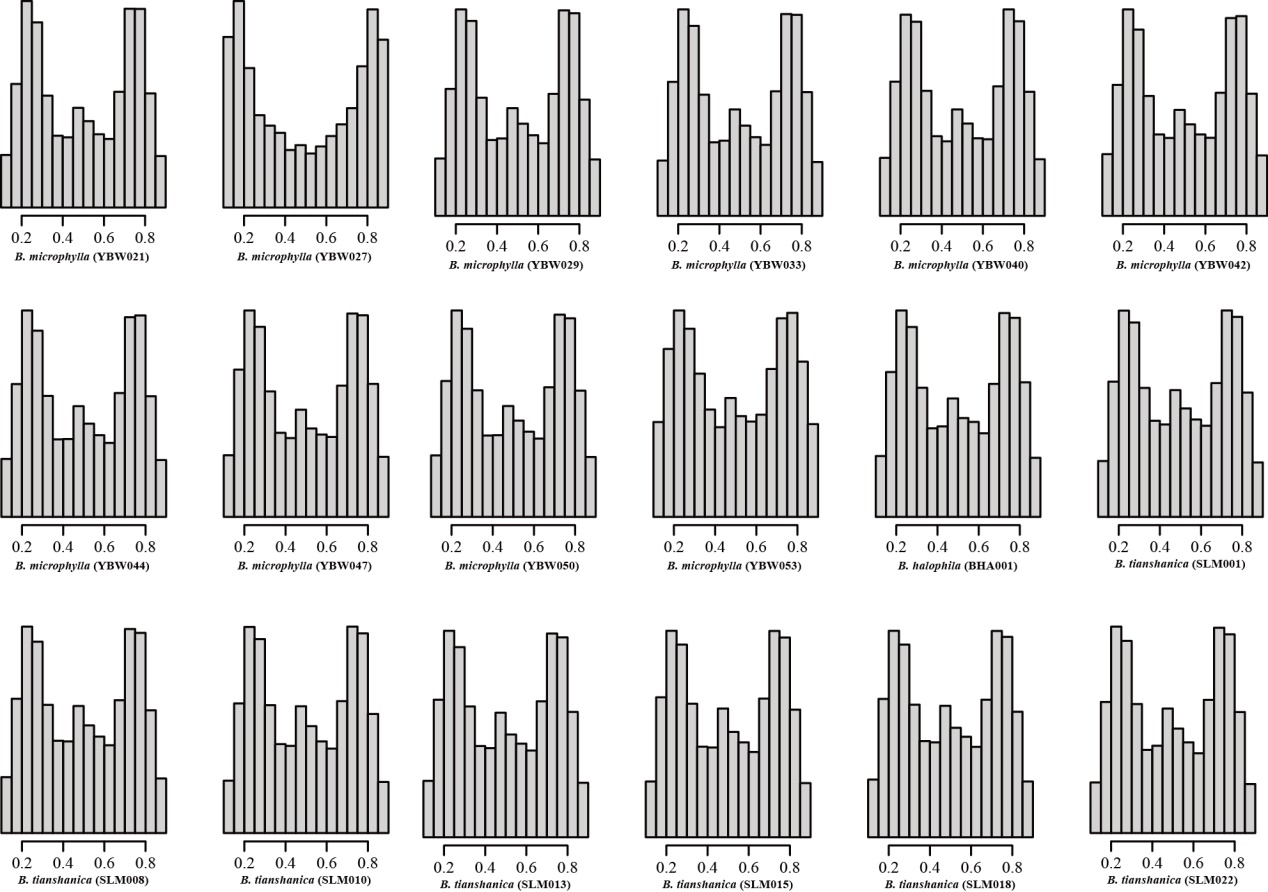

**Figure S5** Phylogenetic tree from the maximum-likelihood analysis of *B. halophila*, *B. microphylla* and *B. tianshanica* using the supermatrix approach based on data from 1,384,918 SNPs. The scale bar below indicates the mean number of nucleotide substitutions per site. Bootstrap percentages of 100 % are not shown. The colored indicates the sequenced samples in the present study.

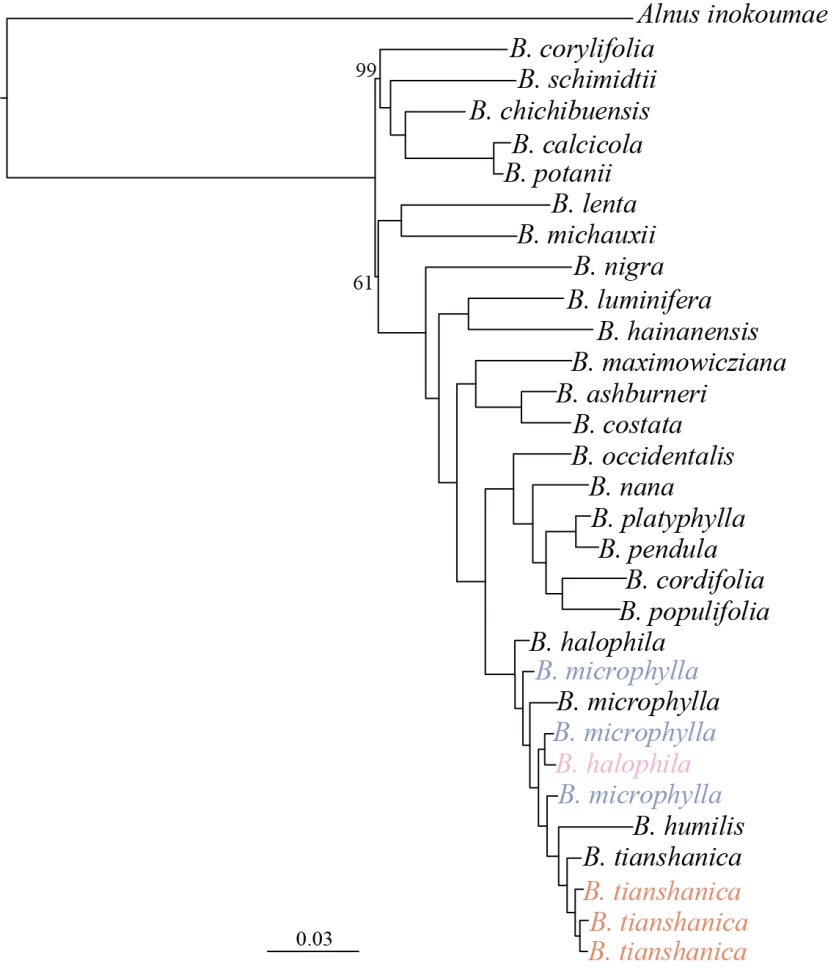
